# Distinct target-specific mechanisms homeostatically stabilize transmission at pre-and post-synaptic compartments

**DOI:** 10.1101/2020.04.07.029108

**Authors:** Pragya Goel, Samantha Nishimura, Karthik Chetlapalli, Xiling Li, Catherine Chen, Dion Dickman

**Author notes:** Correspondence: Dion Dickman, PhD, Department of Neurobiology, University of Southern California, Los Angeles, CA 90089.

## Abstract

Neurons must establish and stabilize connections made with diverse targets, each with distinct demands and functional characteristics. At *Drosophila* neuromuscular junctions, synaptic strength remains stable in a manipulation that simultaneously induces hypo-innervation on one target and hyper-innervation on the other. However, the expression mechanisms that achieve this exquisite target-specific homeostatic control remain enigmatic. Here, we identify the distinct target-specific homeostatic expression mechanisms. On the hypo-innervated target, an increase in postsynaptic glutamate receptor (GluR) abundance is sufficient to compensate for reduced innervation, without any apparent presynaptic adaptations. In contrast, a target-specific reduction in presynaptic neurotransmitter release probability is reflected by a decrease in active zone components restricted to terminals of hyper-innervated targets. Finally, loss of postsynaptic GluRs on one target induces a compartmentalized, homeostatic enhancement of presynaptic neurotransmitter release called presynaptic homeostatic potentiation that can be precisely balanced with the adaptations required for both hypo- and hyper-innervation to maintain stable synaptic strength. Thus, distinct anterograde and retrograde signaling systems operate at pre- and post-synaptic compartments to enable target-specific, homeostatic control of neurotransmission.

## INTRODUCTION

Synapses are spectacularly diverse in their morphology, physiology, and functional characteristics. These differences are reflected in the molecular composition and abundance of synaptic components at heterogenous synaptic subtypes in central and peripheral nervous systems (Atwood and Karunanithi, 2002; Branco and Staras, 2009; O’Rourke et al., 2012). Interestingly, the structure and function of synapses can also vary substantially across terminals of an individual neuron (Fekete et al., 2019; Grillo et al., 2018; Guerrero et al., 2005) and drive input-specific presynaptic plasticity (Letellier et al., 2019). Both Hebbian and homeostatic plasticity mechanisms can work locally and globally at specific synapses to tune synapse function, enabling stable yet flexible ranges of synaptic strength (Diering and Huganir, 2018; Turrigiano, 2012; Vitureira and Goda, 2013). For example, homeostatic receptor scaling globally adjusts GluR abundance, subtype, and/or functionality at dendrites (Turrigiano and Nelson, 2004) yet there is also evidence for synapse specificity (Béïque et al., 2011; Hou et al., 2008; Sutton et al., 2006). Although a number of studies have begun to elucidate the factors that enable both local and global modes of synaptic plasticity at synaptic compartments, it is less appreciated how and why specific synapses undergo plasticity within the context and needs of information transfer in a neural circuit.

One major force that sculpts the heterogeneity of synaptic strength is imposed through the specific targets being innervated. For example, studies at neuromuscular synapses in the stomatogastric system of lobsters have demonstrated that presynaptic terminals of the same motor axon can concurrently undergo facilitation and depression due to differences in the synapses made onto two postsynaptic muscle fibers (Katz et al., 1993). Furthermore, at vertebrate neuromuscular junctions (NMJs), secreted factors from muscles can dictate which motor neurons survive during development and in many cases their neurotransmitter phenotype (Calderó et al., 1998; Schotzinger and Landis, 1990), and a parallel target-dependent control of neuropeptide identity has been shown in the *Drosophila* central nervous system (Allan et al., 2003; Allan and Thor, 2015). In mammalian central neurons, factors such as BDNF secreted from postsynaptic dendrites not only promote neuronal survival but also can homeostatically enhance presynaptic neurotransmitter release and functional properties of neural circuits (Jakawich et al., 2010; Park and Poo, 2013), while postsynaptic signaling through N-Cadherins and mTORC1 can regulate presynaptic function (Henry et al., 2012; Vitureira et al., 2011). Finally, at the *Drosophila* NMJ, presynaptic homeostatic plasticity can be expressed at a subset of terminals on a single motor neuron depending on GluR functionality at particular targets (Li et al., 2018a), demonstrating that this form of homeostatic plasticity is target-specific and strongly suggesting it is also synapse-specific. Together, these studies and others have demonstrated that the physiologic, metabolic, and/or structural properties at terminals of a single neuron can be selectively modulated according to the identity and needs of the targets and synapses they innervate. However, the nature of the trans-synaptic dialogue and the molecular mechanisms that achieve target-specific plasticity are not well understood.

A seminal study published over 20 years ago found that distinct target-specific modulations in synaptic activity maintain stable neurotransmission following biased innervation at terminals of motor neurons at the *Drosophila* NMJ (Davis and Goodman, 1998). In this manipulation, biased innervation is achieved by overexpression of the trans-synaptic cell adhesion factor *fasciculin II* (*fasII*) on one of the two muscle targets innervated by motor neurons (Davis and Goodman, 1998). This leads to hyper-innervation of the target overexpressing *fasII* at the expense of the adjacent target, which is hypo-innervated. Remarkably, synaptic strength, as assessed by electrophysiological recordings, was maintained at levels similar in amplitude to normally innervated NMJ targets. Since this pioneering study, however, the molecular and cellular expression mechanisms that achieve this target-specific homeostatic modulation have remained enigmatic.

We have investigated how terminals of an individual neuron adapt to simultaneous hypo- and hyper-innervation to maintain stable synaptic strength on two adjacent targets. Our analysis reveals that a novel homeostatic signaling system operates in the hypo-innervated target to precisely enhance the abundance of postsynaptic GluRs, offsetting reduced presynaptic neurotransmitter release and stabilizing synaptic strength. In contrast, no apparent adaptations are observed in the hyper-innervated target. Rather, presynaptic release probability is homeostatically reduced, accompanied by a target-specific decrease in the abundance and density of active zone components. Finally, we find that presynaptic homeostatic potentiation can be selectively induced and expressed at synapses on one target and balanced with biased innervation to sustain stable synaptic strength. This work reveals the striking interplay of target-specific homeostats modulating the efficacy of neurotransmission across synaptic terminals.

## RESULTS

### Target-specific mechanisms maintain stable synaptic strength at hypo- and hyper-innervated NMJs

We first sought to reproduce and confirm the biased innervation and synaptic electrophysiology reported in Davis and Goodman, 1998. At *Drosophila* larval NMJs, motor neurons distribute their synaptic terminals roughly evenly between two distinct targets – as demonstrated by the NMJs made onto muscles 6 and 7 (Fig. 1A; left). This stereotyped pattern of innervation can be visualized by immunostaining the NMJ with antibodies that recognize the neuronal membrane (HRP) and synaptic vesicles (Synapsin; SYN), which demonstrates ∼60% boutons on the larger muscle 6 and ∼40% on the smaller muscle 7 (Fig. 1B; left and Table S2). To bias innervation on these targets, we used the *H94-Gal4* driver to drive expression of the cell adhesion molecule *fasciculin II* (*fasII*) early in development selectively on muscle 6 [(M6>FasII; (Davis and Goodman, 1998)]. Immunostaining of M6>FasII NMJs confirmed biased innervation with ∼150% of boutons above controls on muscle 6 (hyper-innervated), and a parallel reduction of ∼50% in boutons on muscle 7 (hypo-innervated) (Fig. 1A-E), consistent with the previous study (Davis and Goodman, 1998). However, despite these opposing changes in bouton numbers, electrophysiological recordings of M6>FasII found that synaptic strength, measured by the excitatory postsynaptic potential (EPSP) amplitude, was similar on both targets and unchanged from their respective controls (Fig. 1C,D,E). This implies target-specific mechanisms modulate neurotransmission on hypo- and hyper-innervated terminals to maintain stable NMJ strength.

**Figure 1:**
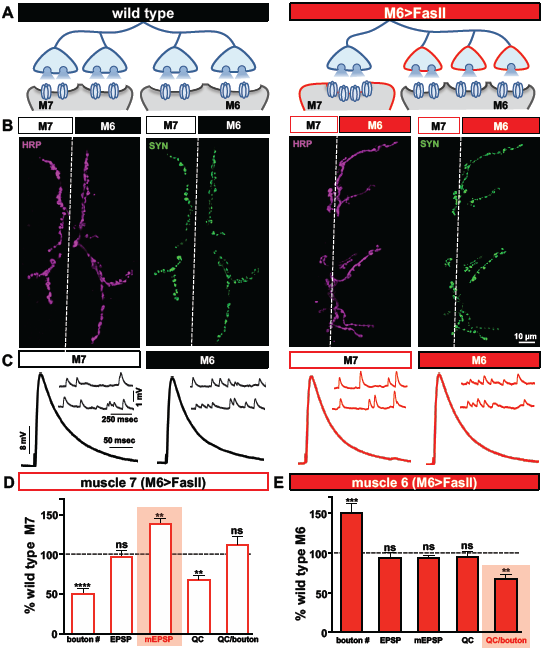
Biased innervation at the NMJ elicits distinct target-specific homeostatic adaptations. **(A)** Schematic of a motor neuron innervating both muscle 6 and 7 at the *Drosophila* larval NMJ. Biased innervation is achieved by overexpressing the cell adhesion factor *fasII* specifically on muscle 6 using *H94-Gal4* (M6>FasII: *w;UAS-FasII/+;H94-Gal4/+*). Red outlines highlight the likely synaptic compartment in which the adaptation occurs. **(B)** Representative images of muscle 6/7 NMJs immunostained with antibodies that recognize the neuronal membrane (HRP) and synaptic vesicles (Synapsin; SYN) in wild type (*w*^*1118*^) and M6>FasII. Note that while boutons labeled by SYN puncta are roughly equally split between muscles 6 and 7 in wild type, M6>FasII causes biased innervation on muscle 6 at the expense of muscle 7. **(C)** Representative electrophysiological traces of recordings from muscles 7 and 6 in wild type and M6>FasII NMJs. Note that while EPSP amplitudes are similar across all muscles, mEPSPs are increased only on muscle 7 of M6>FasII. **(D)** Quantification of bouton number, EPSP amplitude, mEPSP amplitude, quantal content and quantal content normalized per bouton on muscle 7 in M6>FasII. All values are normalized to the values at wild type muscle 7. Enhanced mEPSP amplitude (shaded bar) implies reduced quantal content and no change in quanta released per bouton. **(E)** Quantification of all values in (D) on muscle 6 of M6>FasII normalized to wild type muscle 6 values. Note that estimated quantal content per bouton (shaded bar) is significantly reduced. Error bars indicate ±SEM (n≥16; one-way ANOVA; Table S2). **p<0.01; ***p<0.001; ****p<0.0001; ns=not significant.

To gain insight into how EPSP amplitudes remain similar to baseline values at NMJs with biased innervation, we next examined miniature neurotransmission. On hypo-innervated muscle 7, mEPSP amplitudes were significantly increased by ∼40% compared to baseline values (Fig. 1C,D), as previously observed (Davis and Goodman, 1998). Quantal content (QC) was thus decreased by ∼40%, a value similar in magnitude to the reduction in bouton number (Fig. 1D). In contrast, mEPSP amplitude was not significantly different on the hyper-innervated muscle 6 NMJ compared to baseline (Fig. 1C,E), with no apparent change in quantal content (Fig. 1E), as previously observed (Davis and Goodman, 1998). Finally, analysis of quantal content normalized per bouton on muscle 6 NMJs revealed an ∼30% reduction (Fig. 1E), suggesting a target-specific, homeostatic decrease in presynaptic neurotransmitter release. Together, this data indicates that distinct target-specific mechanisms operate to stabilize neurotransmission at hypo-vs. hyper-innervated NMJs.

### A homeostatic increase in postsynaptic GluR abundance stabilizes synaptic strength on hypo-innervated targets

It was previously reported that at hypo-innervated NMJs following M6>FasII, levels of the postsynaptic GluR subunit GluRIIA were increased (Goel and Dickman, 2018). We considered four additional possible mechanisms that could, in principle, contribute to the observed enhancement in quantal size at hypo-innervated NMJs. First, increased presynaptic vesicle size could lead to enhanced glutamate emitted per vesicle, as has been documented in endocytosis mutants and following overexpression of the vesicular glutamate transporter (Daniels et al., 2004; Goel et al., 2019a). Second, multivesicular release has been observed in some systems (Rudolph et al., 2015) and was raised as a possibility in the original study to potentially explain the increased quantal size (Davis and Goodman, 1998), although there remains no evidence for multi-vesicular release at the fly NMJ. In addition to these two presynaptic mechanisms, other postsynaptic mechanisms are possible that could explain the increased mEPSP amplitude on hypo-innervated NMJs, parallel to the ones that have been documented in mammalian forms of homeostatic receptor scaling (Diering and Huganir, 2018; Turrigiano, 2008). These include increases in the abundance, subtype, and/or functionality of additional postsynaptic GluRs, including GluRIIB-containing receptors, as enhanced levels of GluRIIA-containing GluRs were recently reported at hypo-innervated NMJs in Drosophila (Goel and Dickman, 2018; Goel et al., 2019b).

We therefore examined postsynaptic GluR levels in hypo-innervated targets induced by M6>FasII. At the *Drosophila* NMJ, the postsynaptic response to glutamate is mediated by two subtypes of GluRs, GluRIIA- and GluRIIB-containing receptors. Both subtypes are composed of the essential subunits GluRIIC, GluRIID, and GluRIIE but differ in containing either GluRIIA or GluRIIB subunits (DiAntonio, 2006; Qin et al., 2005). We immunostained hypo-innervated NMJs using antibodies against GluRIIA, GluRIIB, and the common GluRIID subunits and observed an ∼40% decrease in the number of GluR puncta comparted to wild type muscle 7 (Fig. S1A,C), reflecting reduced innervation. However, we found an increase in the intensity of all GluR subunits in hypo-innervated NMJs compared to wild type muscle 7 (Fig. S1B,D). In principle, the 40% increase in GluR abundance is sufficient to explain the increased quantal size and to offset the 40% reduction in quantal content to homeostatically maintain stable synaptic strength despite reduced innervation. Consistent with this, we observed no adaptations in the anatomical or functional number of release sites, nor in the size of the readily releasable pool (Fig. S2). These lines of evidence indicate that presynaptic terminals of hypo-innervated NMJs function similarly to wild type, with presynaptic neurotransmitter released onto the muscle 7 NMJ simply reduced by 40%. Thus, a 40% increase in postsynaptic GluR abundance is sufficient to maintain synaptic strength at hypo-innervated NMJs without reason to invoke other homeostatic adaptations.

### Hyper-innervation induces a homeostatic decrease in presynaptic release probability

We next sought to characterize the expression mechanism that enables stable neurotransmitter output on the hyper-innervated target. It was previously demonstrated that a homeostatic reduction in presynaptic release probability was expressed at hyper-innervated NMJs (Davis and Goodman, 1998). Consistent with this conclusion, and in contrast to the adjacent hypo-innervated NMJs, we did not observe any significant changes in postsynaptic GluR levels (Fig. S1E-H). We next performed a series of electrophysiological assays to probe presynaptic function on the hyper-innervated NMJ. First, we used failure analysis to assess presynaptic release independently of miniature transmission by measuring the number of failed release events in very low extracellular Ca^2+^ concentrations (0.15 mM; see Methods). We observed no significant difference in the failure rates on hyper-innervated NMJs compared to wild type (Fig. 2B), consistent with overall quantal content released being unchanged at these NMJs. Next, we probed release probability by determining paired pulse ratios in moderate and high extracellular Ca^2+^. At 0.4 mM Ca^2+^, we observed an increase in paired pulse facilitation (PPF) at hyper-innervated NMJs compared to wild type (Fig. 2C,D), while at 1.5 mM Ca^2+^, paired-pulse depression (PPD) was reduced at hyper-innervated NMJs (Fig. 2E,F). Since enhanced PPF and reduced PPD is indicative of reduced release probability (Regehr, 2012), these results suggest that a target-specific, homeostatic decrease in presynaptic release probability stabilizes transmission at hyper-innervated NMJs. Although the PPF/PPD recordings suggested reduced release probability at hyper-innervated terminals, the magnitude of the observed decrease (∼25%) was not sufficient to fully compensate for the ∼50% increase in bouton numbers. We found no change in the size of the RRP on hyper-innervated NMJs compared to wild type (Fig. 2G,H), suggesting that the size of the RRP at individual boutons might be reduced on muscle 6 of M6>FasII NMJs. Finally, no change in the total number of functional release sites was observed on hyper-innervated targets (Fig. 2I-K), indicating a reduction in the number of release sites participating in neurotransmission per bouton at hyper-innervated NMJs. Thus, a homeostatic adjustment in the release probability per bouton selectively modulates transmission at hyper-innervated NMJs without measurably impacting release at adjacent hypo-innervated terminals.

**Figure 2:**
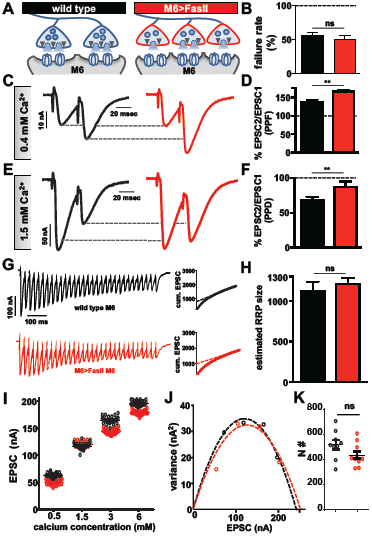
Hyper-innervation elicits a homeostatic decrease in presynaptic release probability. **(A)** Schematic illustrating a reduction in RRP size and functional release site number on hyper-innervated muscle 6. **(B)** Failure analysis reveals no significant change in failure rate on muscle 6 in M6>FasII, consistent with reduced release probability per bouton. **(C)** Representative paired-pulse EPSC traces at 0.4 mM extracellular Ca^2+^ with an interstimulus interval of 16.7 ms in the indicated genotypes. Increased paired-pulse facilitation (PPF) was observed on hyper-innervated targets, consistent with reduced release probability. **(D)** Quantification of PPF ratio (EPSC2/EPSC1). **(E)** Representative paired-pulse EPSC traces at 1.5 mM extracellular Ca^2+^ with an interstimulus interval of 16.7 ms in the indicated genotypes. Enhanced paired-pulse depression (PPD) was observed on hyper-innervated targets, consistent with reduced probability of release. **(F)** Quantification of PPD ratio (EPSC2/EPSC1). **(G)** Representative EPSC recordings of 30 stimuli at 3 mM extracellular Ca^2+^ during a 60 Hz stimulus train in the indicated genotypes. Insets represent the average cumulative EPSC plotted as a function of time. A line fit to the 18^th^-30^th^ stimuli was back-extrapolated to time 0. **(H)** Estimated size of the RRP is unchanged on muscle 6 in M6>FasII compared with wild type, suggesting reduced RRP per bouton. **(I)** Scatter plot EPSC distribution of recordings on muscle 6 from wild type and M6>FasII in the indicated extracellular Ca^2+^ concentrations. **(J)** Variance-mean plots for the indicated genotypes. Dashed lines are the best fit parabolas to the data points. **(K)** Estimated number of functional release sites (N#) obtained from the variance-mean plots in (J) showing no significant difference between the genotypes. Error bars indicate ±SEM (n≥9; one-way ANOVA; Table S2). **p<0.01; ns=not significant.

### A target-specific reduction in both active zone density and intensity is observed at hyper-innervated NMJs

Our electrophysiological data above suggests a reduction in both release probability and the number of functional release sites at individual boutons of hyper-innervated NMJs. In principle, a target-specific reduction in the number and/or function of anatomical release sites could explain these electrophysiological properties. In addition, recent evidence indicates that bi-directional changes in the size and nano-structure of active zone architecture at *Drosophila* NMJs can adjust release probability at individual active zones (Akbergenova et al., 2018; Bohme et al., 2019; Goel et al., 2019a; Gratz et al., 2019). We therefore characterized the number and intensity of individual active zones on hyper-innervated NMJs by immunostaining the central scaffold BRP and endogenously tagged CaV2.1 calcium channels [Cac^sfGFP^; (Gratz et al., 2019)], defining each BRP punctum to be an active zone. Interestingly, while a ∼55% increase in bouton number was observed at hyper-innervated NMJs, the number of active zones was only increased by ∼20%, reflected in a concomitant decrease in active zone density (Fig. 3A-E). Thus, hyper-innervated NMJs exhibit a target-specific reduction in the density of release sites that is sufficient in magnitude to limit the increase in active zones to only about 20% despite an ∼50% increase in innervation.

**Figure 3:**
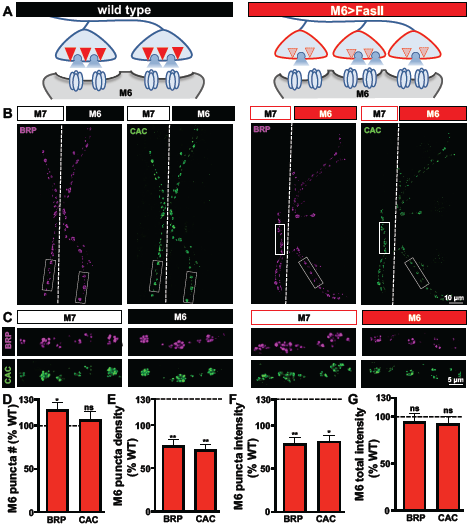
Target-specific reductions in both active zone density and intensity at hyper-innervated NMJs. **(A)** Schematic illustrating a reduction in the number and intensity of active zones at individual boutons on hyper-innervated muscle 6. **(B)** Representative images of muscle 6/7 NMJs in the indicated genotypes (wild type: *cac*^*sfGFP-N*^; M6>FasII: *cac*^*sfGFP-N*^;*UAS-fasII/+;H94-Gal4/+*) immunostained with antibodies against the active zone scaffold *bruchpilot* (BRP) and GFP to label endogenously tagged Ca^2+^ channels (CAC). **(C)** Individual boutons from selected areas (white rectangles) of NMJs stained with BRP and CAC in the indicated genotypes and muscles. Note the reduction in the number and intensity of BRP and CAC puncta specifically on muscle 6 in M6>FasII, while no change is observed on muscle 7 relative to wild type controls. Quantification of BRP and CAC puncta number **(D)** and density **(E)** on muscle 6 in M6>FasII normalized as a percentage of wild type muscle 6 values reveals a small but significant increase in BRP puncta number, while BRP and CAC puncta density is significantly reduced on muscle 6 in M6>FasII. Quantification of BRP and CAC intensity **(F)** shows a significant reduction on muscle 6 in M6>FasII, while the total fluorescence intensity of all BRP and CAC puncta summed across the entire muscle 6 NMJ **(G)** is unchanged compared to wild type muscle 6. Error bars indicate ±SEM (n≥13; one-way ANOVA; Table S2). *p<0.05; **p<0.01; ns=not significant.

We also quantified the intensity of individual BRP puncta on hyper-innervated NMJs and observed an ∼20% decrease in the sum intensity compared to wild type (Fig. 3B,C,F). Similar results for puncta density and intensity were found for Cac^sfGFP^ (Fig. 3B-F). Finally, given these reductions in the density and intensity of active zone components, the total intensity of both BRP and Cac^sfGFP^ per hyper-innervated NMJ was not significantly different from wild type despite the increase in their total number (Fig. 3G). These results parallel recent studies that have shown that while the number and intensity of individual active zones can vary at NMJs, the total abundance of active zone protein remains constant (Goel et al., 2019a; Goel et al., 2019b; Graf et al., 2009). Together, hyper-innervated NMJs express a target-specific reduction in both the number and intensity of release sites and a parallel reduction in presynaptic release probability that stabilizes synaptic strength, while no reciprocal changes are observed at hypo-innervated counterparts.

### Distinct target-specific adaptations can homeostatically balance hyper-innervation and GluR perturbation

When biased innervation of the NMJ is induced through M6>FasII, the hypo-innervated target responds by homeostatically increasing GluR abundance, while the subset of motor neuron terminals that hyper-innervate the adjacent target selectively reduce the number and apparent abundance of active zone components. In our final set of experiments, we sought to determine whether the target-specific homeostatic adaptations triggered by biased innervation could be balanced with an additional target-specific homeostatic challenge. Presynaptic homeostatic potentiation (PHP) is a well-studied form of homeostatic plasticity at the *Drosophila* NMJ. Here, rapid pharmacological or chronic genetic manipulations that diminish postsynaptic GluR functionality trigger a trans-synaptic retrograde signaling system that homeostatically increases presynaptic glutamate release to maintain stable synaptic strength (Frank et al., 2020). Recently, it was demonstrated that GluR knockdown specifically on muscle 6 can trigger PHP selectively at the subset of synapses innervating muscle 6 without influencing transmission at the synaptic terminals of the same motor neuron that innervate the adjacent muscle 7 (Li et al., 2018a), demonstrating a remarkable degree of compartmentalized expression of PHP. We combined these manipulations to induce a simultaneous challenge of biased innervation and GluR loss using *fasII* overexpression combined with *GluRIIA* knockdown selectively on muscle 6 (M6>FasII+GluRIIA^RNAi^). We first validated if the combined manipulation was successful by assaying synaptic growth and GluRIIA levels. Indeed, we observed the expected increase and decrease in bouton numbers on muscles 6 and 7 respectively, with a near absence of GluRIIA immunostaining selectively on muscle 6 (Fig. 4B-F). Thus, target-specific, homeostatic challenges of biased innervation and GluR loss can be simultaneously induced by overexpressing *fasII* and GluRIIA^RNAi^ selectively on muscle 6.

**Figure 4:**
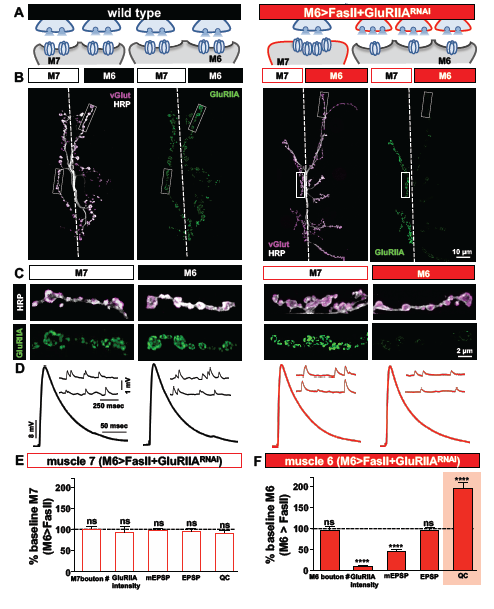
Distinct target-specific adaptations balance hyper-innervation and GluR loss. **(A)** Schematic illustrating the dual manipulation used to both bias innervation and inhibit *GluRIIA* expression specifically on muscle 6 (M6>FasII+GluRIIA^RNAi^: *w;Tub-FRT-STOP-FRT-Gal4,UAS-FLP,UAS-CD8-GFP;H94-Gal4,nSyb-Gal80;/UAS-FasII;UAS-GluRIIA*^*RNAi*^). **(B)** Representative images of muscle 6/7 NMJs from the indicated genotypes immunostained with anti-HRP and -GluRIIA. **(C)** Individual boutons from selected areas (white rectangles) of NMJs shown in (B). Note the enhanced GluRIIA levels on hypo-innervated muscle 7 with loss on hyper-innervated muscle 6. **(D)** Electrophysiological traces of recordings from muscles 7 and 6 in the indicated genotypes. EPSP amplitudes on muscle 7 and 6 in M6>FasII+GluRIIA^RNAi^ are similar to wild-type values. **(E)** Quantification of bouton numbers, GluRIIA puncta intensity, mEPSP amplitude, EPSP amplitude, and quantal content on muscle 7 in M6>FasII+GluRIIA^RNAi^. All values are normalized to baseline (M6>FasII muscle 7); no significant differences are observed. **(F)** Quantification of all values in (D) on muscle 6 of M6>FasII+GluRIIA^RNAi^ normalized to baseline values. Note that while GluRIIA levels and mEPSP amplitudes are significantly reduced, EPSP amplitude remains unchanged because of a homeostatic increase in quantal content, indicating PHP expression. Error bars indicate ±SEM (n≥8; Student’s t-test; Table S2). ****p<0.0001; ns=not significant.

We next performed synaptic electrophysiology at both targets. On the hypo-innervated muscle 7 of M6>FasII+GluRIIA^RNAi^, neurotransmission was indistinguishable from M6>FasII alone, with elevated mEPSP amplitudes, stable EPSP amplitudes, and reduced quantal content observed (Fig. 4D,E). In contrast, on the hyper-innervated muscle 6 of M6>FasII+GluRIIA^RNAi^, mEPSP amplitudes were selectively reduced due to GluR knockdown, but synaptic strength was maintained at baseline levels due to a homeostatic increase in quantal content (Fig. 4D,F). This demonstrates that PHP can be robustly expressed and balanced with the adaptations necessary to adjust release for hyper-innervation in a target-specific manner, without any apparent changes in transmission at adjacent synapses of the hypo-innervated muscle 7. Finally, we tested whether PHP can be acutely induced and balanced at hypo-innervated NMJs after the adjustments made at muscle 6 of M6>FasII+GluRIIA^RNAi^. We applied sub-blocking concentrations of the GluR venom philanthotoxin-433 (PhTx) at NMJs for 10 mins. This acutely induced PHP at wild type NMJs, with reduced mEPSP amplitude but EPSP amplitudes unchanged from baseline due to a rapid, homeostatic increase in quantal content (Fig. S3A-D). Application of PhTx to M6>FasII+ GluRIIA^RNAi^ NMJs had no significant change in mEPSP amplitude or quantal content at muscle 6 due to GluRIIA knockdown (Fig. S3A,B,D). However, PhTx application also induced robust PHP at muscle 7 NMJs in M6>FasII+ GluRIIA^RNAi^, with a significant reduction in mEPSP amplitude but normal EPSP amplitude due to enhanced quantal content (Fig. S3A,C,E). These results demonstrate that presynaptic release sites at terminals of the same neuron can be selectively modulated with exquisite target specificity to compensate for GluR loss and can be superimposed with the homeostatic plasticity induced by biased innervation.

## DISCUSSION

Recent studies have shed light on how neurotransmission is stabilized when synaptic growth and function is challenged (Davis and Muller, 2015; Frank et al., 2020; Goel et al., 2019a; Goel et al., 2019b; Li et al., 2018b). However, less is known about how this stability is maintained when neuronal terminals confront diverse and even opposing challenges in synaptic growth and function. Here, we have combined biased innervation with acute and chronic challenges to postsynaptic GluR function at targets shared by individual neurons. These experiments have revealed two distinct target-specific mechanisms that enable stable transmission despite biased innervation, operating at either pre- or postsynaptic compartments, and that can be balanced with postsynaptic GluR perturbation. Importantly, these processes occur independently, without impacting transmission within the same neuron on neighboring synapses made on the adjacent target. This demonstrates a remarkable degree of compartmentalized autonomy in homeostatic signaling and suggests an independence of local and global homeostats that work in concert to balance synaptic strength.

### Target-specific homeostatic scaling of postsynaptic GluR receptors

We took advantage of a previously established manipulation to bias synaptic innervation using target-specific expression of the trans-synaptic cell adhesion protein FasII (Davis and Goodman, 1998). On the hypo-innervated target, a selective upregulation in postsynaptic GluR abundance was elicited that was sufficient in magnitude to offset reduced neurotransmitter release and stabilize synaptic strength. This scaling of GluR abundance parallels a well-established mechanism of homeostatic synaptic plasticity in mammalian neurons termed *homeostatic receptor scaling* (Chowdhury and Hell, 2018; Diering and Huganir, 2018; Turrigiano, 2008). Although an enhanced upregulation of GluRs occurs in response to reduced activity in mammalian central neurons, the GluR scaling revealed at the fly NMJ is unique. GluRs at the *Drosophila* NMJ are quite stable, in contrast to the rapid dynamics of GluR trafficking observed in dendritic spines of mammalian central neurons. This receptor stasis may reflect a fundamental property of NMJs, where postsynaptic receptors have half-lives of ∼7 days in rodents (Salpeter and Harris, 1983) and over 24 hours in flies (Rasse et al., 2005). Although NMJ receptors appear to be relatively stable under basal conditions and even in mutants in which synaptic transmission and growth are perturbed (Goel et al., 2019b; Lee et al., 2013; Saitoe et al., 2001), there is emerging evidence that specific challenges, including injury and disease, can provoke relatively rapid remodeling of neurotransmitter receptors at postsynaptic compartments of the NMJ (Goel and Dickman, 2018; Palma et al., 2011; Perry et al., 2017; Rich and Lichtman, 1989). The temporal regulation and dynamics of the GluR plasticity we have characterized at the NMJ in response to hypo-innervation is unclear but likely to be intertwined with the developmental processes of NMJs as well as synaptic growth.

The induction mechanism involved in how reduced innervation is sensed to ultimately instruct an adaptive increase in postsynaptic GluR abundance is unclear. It is notable that while hypo-innervation in the M6>FasII manipulation elicits GluR scaling, a variety of mutations that lead to synaptic undergrowth do not consistently change receptor levels (Banovic et al., 2010; Goel et al., 2019b; Kaufmann et al., 2002; Marqués et al., 2002). Further, mutations that severely reduce neurotransmitter release, including *synaptotagmin* and *complexin* mutants, do not change GluR levels (Huntwork and Littleton, 2007; Lee et al., 2013; Saitoe et al., 2001). Hence, hypo-innervation and/or reduced neurotransmitter release alone is unlikely to be sufficient to induce postsynaptic GluR scaling. Rather, this form of homeostatic plasticity may be dependent on the phenomenon of biased innervation between two targets shared by a single neuron itself, implying some signaling between the motor neuron and/or the adjacent muscles are involved. What is clear is that the postsynaptic signal transduction system that mediates hypo-innervation-dependent GluR scaling is distinct from that which mediates retrograde PHP signaling, as GluR scaling can still be expressed in conditions in which postsynaptic PHP signaling is blocked (Goel and Dickman, 2018). Finally, the capacity for PHP to be robustly expressed at NMJs that have undergone GluR scaling illuminates postsynaptic GluRs to be central targets for both the induction and expression of homeostatic synaptic plasticity.

### Target-specific modulation of active zones

In contrast to the exclusively postsynaptic adaptation observed in response to reduced innervation, an entirely presynaptic mechanism stabilizes synaptic strength at hyper-innervated muscles, expressed by a target-specific reduction in the number and intensity of active zone components. Although a similar reduction in the abundance of active zone proteins at individual release sites has recently been found in mutations that cause synaptic overgrowth at the NMJ (Goel et al., 2019a; Goel et al., 2019b), the adaptations observed in the case of hyper-innervation is distinct in that they are 1) target-specific and 2) involve a reduction in active zone number in addition to protein abundance. It is remarkable that both the number and abundance of active zone components can be selectively reduced and calibrated at hyper-innervated terminals without any apparent changes at adjacent terminals shared by the same neuron on the hypo-innervated target. One attractive pathway that could be controlled to subserve this target-specific modulation of active zone structure might involve the lysosomal adaptor Arl-8. Arl-8 regulates the delivery of synaptic vesicle and active zone cargo to synapses (Klassen et al., 2010; Vukoja et al., 2018), and was recently shown to promote the delivery of synaptic cargo necessary to remodel active zones during PHP (Goel et al., 2019a). Because the abundance of active zone components are remodeled during PHP (Bohme et al., 2016; Goel et al., 2017; Gratz et al., 2019; Weyhersmuller et al., 2011) through an *arl-8* dependent mechanism (Goel et al., 2019a), and PHP can be expressed at a subset of terminals with target-specificity (Li et al., 2018a), it is tempting to speculate that Arl-8 may also be involved in the target-specific reduction in active zones following hyper-innervation.

### Biased innervation, presynaptic homeostatic plasticity, and information transfer at synapses

Global synaptic strength is established during development through intrinsic genetic programs and a dialogue between pre- and post-synaptic compartments. Robustness in this process is ensured by signaling systems that have the capacity to sense and adapt to deviations outside of physiological ranges, such as reductions or enhancements in synaptic growth (Goel et al., 2019b; Goel et al., 2019c; Keck et al., 2013; Tripodi et al., 2008; Yuan et al., 2011). Superimposed on this foundation are forms of plasticity such as PHP, which appear to operate as independent homeostats to maintain stable information transfer at synapses and within neural circuits. Presynaptic terminals of a neuron therefore do not function as unitary computational units but are rather compartmentally specialized and flexible according to the physiologic needs of their targets during development and also following homeostatic challenges. In addition to this target-specificity, there is also evidence for input-specificity across dendrites that have the capacity to homeostatically modulate strength in rodent hippocampal neurons (Jia et al., 2010; Katz et al., 2009; Letellier et al., 2019; Stuart and Spruston, 2015). This remarkable control of synaptic activity enables the flexibility to locally adjust synaptic strengths through input- and target-specificity while stabilizing overall network activity and information processing.

## MATERIALS AND METHODS

### Fly stocks

*Drosophila* stocks were raised at 25°C on standard molasses food. The *w*^*1118*^ strain is used as the wild type control unless otherwise noted as this is the genetic background in which all genotypes are bred. Details of all stocks and their sources are listed in the Reagents and Resource Table S1.

### Immunocytochemistry

Third-instar larvae were dissected in ice cold 0 Ca^2+^ HL-3 and immunostained using a standard protocol as described (Perry et al., 2017). In brief, larvae were either fixed in Bouin’s fixative for 5 min (Sigma, HT10132-1L), 100% ice-cold ethanol for 5 min, or 4% paraformaldehyde (PFA) for 10 min. Larvae were then washed with PBS containing 0.1% Triton X-100 (PBST) for 30 min, blocked with 5% Normal Donkey Serum followed by overnight incubation in primary antibodies at 4°C. Preparations were then washed 3x in PBST, incubated in secondary antibodies for 2 hours, washed 3x in PBST, and equilibrated in 70% glycerol. Prior to imaging, samples were mounted in VectaShield (Vector Laboratories). Details of all antibodies, their source, dilution, and references are listed in Table S1.

### Confocal imaging and analysis

Samples were imaged using a Nikon A1R Resonant Scanning Confocal microscope equipped with NIS Elements software and a 60x APO 1.4 NA oil immersion objective using separate channels with four laser lines (405, 488, 561, and 637 nm) at room temperature. Boutons were counted using NMJs stained with anti-Synapsin or -vGlut co-stained with anti-HRP on muscle 6/7 of segment A2 and A3, considering each Synapsin or vGlut punctum to be a bouton. For fluorescence quantifications of postsynaptic GluRs and active zone proteins, all genotypes within a data set were immunostained in the same tube with identical reagents, then mounted and imaged in the same session. Z-stacks were obtained using identical settings for all genotypes with z-axis spacing between 0.15-0.2 μm within an experiment and optimized for detection without saturation of the signal. Maximum intensity projections were used for quantitative image analysis with the NIS Elements General Analysis toolkit.

To quantify *sum puncta intensity*, the total fluorescence intensity signal of individual puncta were calculated without regard to area as described (Goel et al., 2019a). For each particular sample set, thresholds were optimized to capture the dynamic range of intensity levels within the wild type sample. This same threshold was then used to image all other genotypes in the sample set, and all intensities were normalized to wild type values within an experimental set. Active zones too closely spaced to be resolved (∼5% of all analyzed) were excluded from the analysis. Spot detection in the Nikon Elements Software was used to identify individual BRP and Cac puncta as it resolves closely spaced puncta more accurately compared to thresholding. Finally, to quantify *total intensity per muscle*, the sum fluorescence intensity for each individual punctum was summed across the entire muscle. For calculation of BRP and Cac puncta density, the total number of puncta at a particular muscle was divided by the neuronal membrane area labeled by HRP spanning that muscle (Goel et al., 2019a). For image representation only, the gain and contrast were adjusted identically for all genotypes within a dataset. To show representative images of individual boutons, a particular area was selected from the entire NMJ (denoted with a white box) and rotated and cropped to demonstrate changes at boutons more clearly.

### Electrophysiology

All dissections and recordings were performed in modified HL-3 saline (Kikuma et al., 2017; Stewart et al., 1994) containing (in mM): 70 NaCl, 5 KCl, 10 MgCl_2_, 10 NaHCO_3_, 115 Sucrose, 5 Trehelose, 5 HEPES, and 0.4 CaCl_2_ (unless otherwise specified), pH 7.2. Neuromuscular junction sharp electrode (electrode resistance between 10-35 MΩ) recordings were performed on muscles 6 and 7 of abdominal segments A2 and A3 in wandering third-instar larvae (Kiragasi et al., 2017). Recordings were performed on an Olympus BX61 WI microscope using a 40x/0.80 NA water-dipping objective. Recordings were acquired using an Axoclamp 900A amplifier, Digidata 1440A acquisition system and pClamp 10.5 software (Molecular Devices). Electrophysiological sweeps were digitized at 10 kHz and filtered at 1 kHz. Data were analyzed using Clampfit (Molecular devices), MiniAnalysis (Synaptosoft), and Excel (Microsoft) software.

Miniature excitatory postsynaptic potentials (mEPSPs) were recorded in the absence of any stimulation and cut motor axons were stimulated to elicit excitatory postsynaptic potentials (EPSPs). An ISO-Flex stimulus isolator (A.M.P.I.) was used to modulate the amplitude of stimulatory currents. Intensity was adjusted for each cell, set to consistently elicit responses from both neurons innervating the muscle segment, but avoiding overstimulation. Average mEPSP, EPSP, and quantal content were calculated for each genotype. Muscle input resistance (R_in_) and resting membrane potential (V_rest_) were monitored during each experiment. Recordings were rejected if the V_rest_ was more depolarized than -60 mV, if the R_in_ was less than 5 MΩ, or if either measurement deviated by more than 10% during the course of the experiment. Larvae were incubated with or without philanthotoxin-433 (PhTx; Sigma; 20 μM) and resuspended in HL-3 for 10 min as described (Dickman and Davis, 2009; Frank et al., 2006).

The readily releasable pool (RRP) size was estimated by analyzing cumulative EPSC amplitudes while recording using a two-electrode voltage clamp (TEVC) configuration as described (Goel et al., 2019c). Muscles were clamped at -80 mV and EPSCs were evoked with a 60 Hz, 60 stimulus train while recording in 3 mM Ca^2+^ HL-3. A line fit to the linear phase (stimuli # 18–30) of the cumulative EPSC data was back-extrapolated to time 0. The RRP value was estimated by determining the extrapolated EPSC value at time 0 and dividing this value by the average mEPSC amplitude.

Data used in the variance-mean plot was obtained from TEVC recordings using an initial 0.5 mM Ca^2+^ concentration, which was subsequently increased to 1.5, 3.0, and 6.0 mM through saline exchange using a peristaltic pump (Langer Instruments, BT100-2J). EPSC amplitudes were monitored during the exchange, and 30 EPSC (0.5 Hz stimulation rate) events were performed in each calcium condition (Li et al., 2018a). To obtain the variance-mean plot, the variance (squared standard deviation) and mean (averaged evoked amplitude) were calculated from the 30 EPSCs at each individual Ca^2+^ concentration. The variance was then plotted against the mean for each specific calcium condition using MATLAB software (MathWorks, USA). One additional data point, in which variance and mean are both theoretically at 0, was used for Ca^2+^-free saline. Data from these five conditions were fit with a standard parabola (variance = Q*I □-I □^2^/N), where Q is the quantal size, I □ is the mean evoked amplitude (x-axis), and N is the number of functional release sites. N, as a parameter of the standard parabola, was directly calculated for each cell by best parabolic fit.

Failure analysis was performed in HL-3 solution containing 0.15 mM CaCl_2_. At this extracellular Ca^2+^ concentration, approximately half of the stimulations evoked responses in the muscle in wild type larvae. A total of 40 trials (stimulations) were performed at each NMJ in all genotypes. Failure rate was obtained by dividing the total number of failures by the total number of trials (40). Paired-pulse recordings were performed at a Ca^2+^ concentration of 0.3 mM to assay facilitation (PPF) and 1.5 mM for depression (PPD). Following the first stimulation, a second EPSC was evoked at an interstimulus interval of 16.66 msec. Paired-pulse ratios were calculated as the difference between the second peak and the maximum value between both peaks (corresponding to the starting point of the second response) divided by the first amplitude.

### Statistical analysis

Data were analyzed using GraphPad Prism (version 7.0) or Microsoft Excel software (version 16.22). Sample values were tested for normality using the D’Agostino & Pearson omnibus normality test which determined that the assumption of normality of the sample distribution was not violated. Data were then compared using either a one-way ANOVA and tested for significance using a Tukey’s multiple comparison test or using an unpaired 2-tailed Student’s t-test with Welch’s correction. All data are presented as mean +/-SEM; n indicates sample number and p denotes the level of significance assessed as p<0.05 (*), p<0.01 (**), p<0.001 (***), p<0.0001 (****); ns=not significant. Statistics of all experiments are summarized in Table S2.

## ACKNOWLEDGEMENTS

We acknowledge the Developmental Studies Hybridoma Bank (Iowa, USA) for antibodies used in this study, and the Bloomington Drosophila Stock Center for fly stocks (NIH P40OD018537). We thank Giwoo Kim and Sarah Perry for technical contributions at early stages of this project. P.G. was supported in part by a USC Provost Graduate Research Fellowship. This work was supported by grants from the National Institutes of Health (NS111414 and NS091546) to D.D.

## AUTHOR CONTRIBUTIONS

PG and DD designed the research; PG, SN, KC, CC, and XL performed the research; PG analyzed the data; PG and DD wrote the paper.

## FIGURE LEGENDS

**Figure S1: Postsynaptic GluR levels are enhanced at hypo-innervated targets but do not change at hyper-innervated targets. (A)** Schematic illustrating enhanced levels of both GluRIIA- and GluRIIB-containing receptors on hypo-innervated muscle 7 in M6>FasII. **(B)** Representative images of individual boutons at NMJs immunostained with antibodies that recognize the postsynaptic GluR subunits GluRIIA, GluRIIB, and GluRIID. Quantification of the indicated GluR puncta number **(C)** and intensity **(D)** normalized to wild type muscle 7 reveals a significant reduction in GluR puncta number but increase in puncta intensity at hypo-innervated muscle 7. **(E)** Schematic illustrating that no changes in postsynaptic GluR levels are induced at hyper-innervated NMJs. **(F)** Representative images of individual boutons in the indicated genotypes at muscle 6 NMJs immunostained with anti-GluRIIA, -GluRIIB, and -GluRIID. **(G)** Quantification of GluR puncta number normalized to wild type muscle 6 demonstrates a significant increase in M6>FasII. **(H)** The total sum intensity of individual GluRIIA, GluRIIB, and GluRIID puncta on muscle 6 NMJs is unchanged in M6>FasII compared to wild type. Error bars indicate ±SEM (n≥9; one-way ANOVA; Table S2). **p<0.01; ***p<0.001; ns=not significant.

**Figure S2: No apparent changes in presynaptic function and the size, density or intensity of active zones are observed at hypo-innervated NMJs**.

**(A)** Representative images of individual boutons at muscle 7 NMJs in wild type and hypo-innervated M6>FasII immunostained with an antibody against the active zone scaffold BRP. **(B)** Quantification of BRP puncta number, density, and intensity on muscle 7 in M6>FasII normalized to wild-type values. A significant decrease in BRP puncta number but no change in density is observed. No significant change in the total intensity of each individual BRP puncta is observed (puncta sum intensity), while the total fluorescence intensity of all BRP puncta summed across the entire muscle 7 NMJ is reduced in hypo-innervated targets. **(C)** Representative EPSC traces of 30 stimuli at 3 mM extracellular Ca^2+^ during a 60 Hz stimulus train recorded in two-electrode voltage clamp (TEVC) configuration in the indicated genotypes. Graphical insets show the average cumulative EPSC plotted as a function of time. A line fit to the 18^th^-30^th^ stimuli was back-extrapolated to time 0. **(D)** The size of the RRP is reduced by a proportion similar to the reduction in quantal content on muscle 7 in M6>FasII. **(E)** Scatter plot EPSC distribution of recordings from muscle 7 in wild type (black circles) and M6>FasII (red triangles) in the indicated extracellular Ca^2+^ concentrations. **(F)** Variance-mean plots for the indicated genotypes. Variance was plotted against the mean amplitude of 30 EPSCs recorded at 0.2 Hz from the Ca^2+^ concentrations detailed in (D). Lines are the best fit parabolas to the data points. **(G)** Estimated number of functional release sites (N #) obtained from the variance-mean plots in (H) showing a significant reduction on muscle 7 of M6>FasII compared to wild type muscle 7. Error bars indicate ±SEM (n≥9; one-way ANOVA; Table S2). **p<0.01; ***p<0.001; ns=not significant.

**Figure S3: PHP can be rapidly expressed at hypo-innervated targets to balance synaptic strength. (A)** Schematics and representative traces illustrating the acute application of PhTx on wild type and M6>FasII+GluRIIA^RNAi^ NMJs. Note that PhTx application reduces mEPSP amplitudes on hypo-innervated targets, yet EPSP amplitudes remain similar to baseline values because of a homeostatic increase in presynaptic neurotransmitter release, indicating robust PHP expression. **(B**,**C**,**D**,**E)** Quantification of mEPSP amplitude, EPSP amplitude and quantal content values normalized to baseline values (wild type muscle 7 in (B) and 6 in (D); M6>FasII+GluRIIA^RNAi^ muscle 7 (C) or 6 (E)). Error bars indicate ±SEM (n≥8; Student’s t-test; Table S2). ****p<0.0001; ns=not significant.

## REFERENCES

Akbergenova, J., K.L. Cunningham, Y.V. Zhang, S. Weiss, and J.T. Littleton. 2018. Characterization of developmental and molecular factors underlying release heterogeneity at Drosophila synapses. Elife. 7:e38268.

Allan, D.W., S.E. St Pierre, I. Miguel-Aliaga, and S. Thor. 2003. Specification of neuropeptide cell identity by the integration of retrograde BMP signaling and a combinatorial transcription factor code. Cell. 113:73–86.

Allan, D.W., and S. Thor. 2015. Transcriptional selectors, masters, and combinatorial codes: regulatory principles of neural subtype specification. Wiley Interdiscip Rev Dev Biol. 4:505–528.

Atwood, H.L., and S. Karunanithi. 2002. Diversification of synaptic strength: presynaptic elements. Nat Rev Neurosci. 3:497–516.

Banovic, D., O. Khorramshahi, D. Owald, C. Wichmann, T. Riedt, W. Fouquet, R. Tian, S.J. Sigrist, and H. Aberle. 2010. Drosophila neuroligin 1 promotes growth and postsynaptic differentiation at glutamatergic neuromuscular junctions. Neuron. 66:724–738.

Béïque, J.C., Y. Na, D. Kuhl, P.F. Worley, and R.L. Huganir. 2011. Arc-dependent synapse-specific homeostatic plasticity. PNAS. 108:816–821.

Bohme, M.A., C. Beis, S. Reddy-Alla, E. Reynolds, M.M. Mampell, A.T. Grasskamp, J. Lutzkendorf, D.D. Bergeron, J.H. Driller, H. Babikir, F. Gottfert, I.M. Robinson, C.J. O’Kane, S.W. Hell, M.C. Wahl, U. Stelzl, B. Loll, A.M. Walter, and S.J. Sigrist. 2016. Active zone scaffolds differentially accumulate Unc13 isoforms to tune Ca(2+) channel-vesicle coupling. Nat Neurosci. 19:1311–1320.

Bohme, M.A., A.W. McCarthy, A.T. Grasskamp, C.B. Beuschel, P. Goel, B. Jusyte, D. Laber, S. Huang, U. Rey, A.G. Petzold, M. Lehmann, F. Goettfert, P. Haghighi, S.W. Hell, D. Owald, D. Dickman, S.J. Sigrist, and A.M. Walter. 2019. Rapid active zone remodeling consolidates presynaptic potentiation. Nat Commun. 10:1085.

Branco, T., and K. Staras. 2009. The probability of neurotransmitter release: variability and feedback control at single synapses.. Nat Rev Neurosci. 10:373–383.

Calderó, J., D. Prevette, X. Mei, R.A. Oakley, L. Li, C. Milligan, L. Houenou, M. Burek, and R.W. Oppenheim. 1998. Peripheral target regulation of the development and survival of spinal sensory and motor neurons in the chick embryo. J Neurosci. 18:356–370.

Chowdhury, D., and J.W. Hell. 2018. Homeostatic synaptic scaling: molecular regulators of synaptic AMPA-type glutamate receptors. F1000Res. 7:234.

Clements, J.D., and R.A. Silver. 2000. Unveiling synaptic plasticity: a new graphical and analytical approach. Trends Neurosci. 23:105–113.

Daniels, R.W., C.A. Collins, M.V. Gelfand, J. Dant, E.S. Brooks, D.E. Krantz, and A. DiAntonio. 2004. Increased expression of the Drosophila vesicular glutamate transporter leads to excess glutamate release and a compensatory decrease in quantal content. J Neurosci. 24:10466–10474.

Davis, G.W., and C.S. Goodman. 1998. Synapse-specific control of synaptic efficacy at the terminals of a single neuron. Nature. 392:82–86.

Davis, G.W., and M. Muller. 2015. Homeostatic control of presynaptic neurotransmitter release. Annu Rev Physiol. 77:251–270.

DiAntonio, A. 2006. Glutamate receptors at the Drosophila neuromuscular junction. Intl. Rev. Neurobiol. 75:165–179.

DiAntonio, A., S.A. Petersen, M. Heckmann, and C.S. Goodman. 1999. Glutamate receptor expression regulates quantal size and quantal content at the Drosophila neuromuscular junction. J Neurosci. 19:3023–3032.

DiAntonio, A., and T.L. Schwarz. 1994. The effect on synaptic physiology of synaptotagmin mutations in Drosophila. Neuron. 12:909–920.

Dickman, D.K., and G.W. Davis. 2009. The schizophrenia susceptibility gene dysbindin controls synaptic homeostasis. Science. 326:1127–1130.

Diering, G.H., and R.L. Huganir. 2018. The AMPA Receptor Code of Synaptic Plasticity. Neuron. 100:314–329.

Fekete, A., Y. Nakamura, Y.M. Yang, S. Herlitze, M.D. Mark, D.A. DiGregorio, and L.Y. Wang. 2019. Underpinning heterogeneity in synaptic transmission by presynaptic ensembles of distinct morphological modules. Nat Commun. 10:826.

Frank, C.A., T.D. James, and M. Müller. 2020. Homeostatic control of Drosophila neuromuscular junction function. Synapse. 74:e22133.

Frank, C.A., M.J. Kennedy, C.P. Goold, K.W. Marek, and G.W. Davis. 2006. Mechanisms underlying the rapid induction and sustained expression of synaptic homeostasis. Neuron. 52:663–677.

Goel, P., D.D. Bergeron, M. Bohme, L. Nunnelly, M. Lehmann, C. Buser, A.M. Walter, S.J. Sigrist, and D.K. Dickman. 2019a. Homeostatic scaling of active zone scaffolds maintains global synaptic strength. J Cell Biol. 218:1706–1724.

Goel, P., and D. Dickman. 2018. Distinct homeostatic modulations stabilize reduced postsynaptic receptivity in response to presynaptic DLK signaling. Nat Commun. 9:1856.

Goel, P., M. Khan, S. Howard, G. Kim, B. Kiragasi, K. Kikuma, and D. Dickman. 2019b. A Screen for Synaptic Growth Mutants Reveals Mechanisms That Stabilize Synaptic Strength. J Neurosci. 39:4051–4065.

Goel, P., X. Li, and D. Dickman. 2017. Disparate Postsynaptic Induction Mechanisms Ultimately Converge to Drive the Retrograde Enhancement of Presynaptic Efficacy. Cell Rep. 21:2339–2347.

Goel, P., X. Li, and D. Dickman. 2019c. Estimation of the Readily Releasable Synaptic Vesicle Pool at the Drosophila Larval Neuromuscular Junction. Bio Protoc. 9:e3127.

Graf, E.R., R.W. Daniels, R.W. Burgess, T.L. Schwarz, and A. DiAntonio. 2009. Rab3 dynamically controls protein composition at active zones. Neuron. 64:663–677.

Gratz, S.J., P. Goel, J.J. Bruckner, R.X. Hernandez, K. Khateeb, G. Macleod, D. Dickman, and K.M. O’Connor-Giles. 2019. Endogenous tagging reveals differential regulation of Ca2+ channels at single AZs during presynaptic homeostatic potentiation and depression. J Neurosci. 39:2416–2429.

Grillo, F.W., G. Neves, A. Walker, G. Vizcay-Barrena, R.A. Fleck, T. Branco, and J. Burrone. 2018. A Distance-Dependent Distribution of Presynaptic Boutons Tunes Frequency-Dependent Dendritic Integration. Neuron. 99:275–282.

Guerrero, G., D.F. Reiff, G. Agarwal, R.W. Ball, A. Borst, C.S. Goodman, and E.Y. Isacoff. 2005. Heterogeneity in synaptic transmission along a Drosophila larval motor axon. Nat Neurosci. 8:1188–1196.

Han, T.H., P. Dharkar, M.L. Mayer, and M. Serpe. 2015. Functional reconstitution of Drosophila melanogaster NMJ glutamate receptors. PNAS. 112:6182–6187.

Henry, F.E., A.J. McCartney, R. Neely, A.S. Perez, C.J. Carruthers, E.L. Stuenkel, K. Inoki, and M.A. Sutton. 2012. Retrograde changes in presynaptic function driven by dendritic mTORC1. J Neurosci. 32:17128–17142.

Hou, Q., D. Zhang, L. Jarzylo, R.L. Huganir, and H.Y. Man. 2008. Homeostatic regulation of AMPA receptor expression at single hippocampal synapses. PNAS. 105:775–780.

Huntwork, S., and J.T. Littleton. 2007. A complexin fusion clamp regulates spontaneous neurotransmitter release and synaptic growth. Nat Neurosci. 10:1235–1237.

Jakawich, S.K., H.B. Nasser, M.J. Strong, A.J. McCartney, A.S. Perez, N. Rakesh, C.J. Carruthers, and M.A. Sutton. 2010. Local presynaptic activity gates homeostatic changes in presynaptic function driven by dendritic BDNF synthesis. Neuron. 68:1143–1158.

Jia, H., N.L. Rochefort, X. Chen, and A. Konnerth. 2010. Dendritic organization of sensory input to cortical neurons in vivo. Nature. 464:1307–1312.

Katz, P.S., M.D. Kirk, and C.K. Givind. 1993. Facilitation and Depression at Different Branches of the Same Motor Axon: Evidence for Presynaptic Differences in Release. J Neurosci. 13:3075–3089.

Katz, Y., V. Menon, D.A. Nicholson, Y. Geinisman, W.L. Kath, and N. Spruston. 2009. Synapse distribution suggests a two-stage model of dendritic integration in CA1 pyramidal neurons. Neuron. 63:171–177.

Kaufmann, N., J. DeProto, R. Ranjan, H. Wan, and D. Van Vactor. 2002. Drosophila liprin-alpha and the receptor phosphatase Dlar control synapse morphogenesis. Neuron. 34:27–38.

Keck, T., G.B. Keller, R.I. Jacobsen, U.T. Eysel, T. Bonhoeffer, and M. HuLbener. 2013. Synaptic scaling and homeostatic plasticity in the mouse visual cortex in vivo. Neuron. 80:327–334.

Kikuma, K., X. Li, D. Kim, D. Sutter, and D.K. Dickman. 2017. Extended Synaptotagmin Localizes to Presynaptic ER and Promotes Neurotransmission and Synaptic Growth in Drosophila. Genetics. 207:993–1007.

Kiragasi, B., J. Wondolowski, Y. Li, and D.K. Dickman. 2017. A Presynaptic Glutamate Receptor Subunit Confers Robustness to Neurotransmission and Homeostatic Potentiation. Cell Rep. 19:2694–2706.

Klassen, M.P., Y.E. Wu, C.I. Maeder, I. Nakae, J.G. Cueva, E.K. Lehrman, M. Tada, K. Gengyo-Ando, G.J. Wang, M. Goodman, S. Mitani, K. Kontani, T. Katada, and K. Shen. 2010. An Arf-like Small G Protein, ARL-8, Promotes the Axonal Transport of Presynaptic Cargoes by Suppressing Vesicle Aggregation. Neuron. 66:710–723.

Lee, J., Z. Guan, Y. Akbergenova, and J.T. Littleton. 2013. Genetic analysis of synaptotagmin C2 domain specificity in regulating spontaneous and evoked neurotransmitter release. J Neurosci. 33:187–200.

Letellier, M., F. Levet, O. Thoumine, and Y. Goda. 2019. Differential role of pre- and postsynaptic neurons in the activity-dependent control of synaptic strengths across dendrites. PLoS Biol. 17:e2006223.

Li, X., P. Goel, C. Chen, V. Angajala, X. Chen, and D. Dickman. 2018a. Synapse-specific and compartmentalized expression of presynaptic homeostatic potentiation. Elife. 7:e34338.

Li, X., P. Goel, J. Wondolowski, J. Paluch, and D. Dickman. 2018b. A Glutamate Homeostat Controls the Presynaptic Inhibition of Neurotransmitter Release. Cell Rep. 23:1716–1727.

Littleton, J.T., H.J. Bellen, and M.S. Perin. 1993. Expression of synaptotagmin in Drosophila reveals transport and localization of synaptic vesicles to the synapse. Development. 118:1077–1088.

Marqués, G., H. Bao, T.E. Haerry, M.J. Shimell, P. Duchek, B. Zhang, and M.B. O’Connor. 2002. The Drosophila BMP type II receptor Wishful Thinking regulates neuromuscular synapse morphology and function. Neuron. 33:529–543.

O’Rourke, N.A., N.C. Weiler, K.D. Micheva, and S.J. Smith. 2012. Deep Molecular Diversity of Mammalian Synapses: Why It Matters and How to Measure It. Nat Rev Neurosci. 13:365–379.

Palma, E., M. Inghilleri, L. Conti, C. Deflorio, V. Frasca, A. Manteca, F. Pichiorric, C. Roseti, G. Torchia, C. Limatola, F. Grassi, and R. Miledi. 2011. Physiological characterization of human muscle acetylcholine receptors from ALS patients. PNAS. 108:20184–20188.

Park, H., and M.M. Poo. 2013. Neurotrophin regulation of neural circuit development and function. Nat Rev Neurosci. 14:7–23.

Perry, S., Y. Han, A. Das, and D.K. Dickman. 2017. Homeostatic plasticity can be induced and expressed to restore synaptic strength at neuromuscular junctions undergoing ALS-related degeneration. Human Mol Genet. 26:4153–4167.

Qin, G., T. Schwarz, R.J. Kittel, A. Schmid, T.M. Rasse, D. Kappei, E. Ponimaskin, M. Heckmann, and S.J. Sigrist. 2005. Four different subunits are essential for expressing the synaptic glutamate receptor at neuromuscular junctions of Drosophila. J Neurosci. 25:3209–3218.

Rasse, T.M., W. Fouquet, A. Schmid, R.J. Kittel, S. Mertel, C.B. Sigrist, M. Schmidt, A. Guzman, C. Merino, G. Qin, C. Quentin, F.F. Madeo, M. Heckmann, and S.J. Sigrist. 2005. Glutamate receptor dynamics organizing synapse formation in vivo. Nat Neurosci. 8:898–905.

Regehr, W.G. 2012. Short-term presynaptic plasticity. Cold Spring Harb Perspect Biol. 4:a005702.

Rich, M.M., and J.W. Lichtman. 1989. In vivo visualization of pre- and postsynaptic changes during synapse elimination in reinnervated mouse muscle. J Neurosci. 5:1781–1805.

Rosenmund, C., and C.F. Stevens. 1996. Definition of the readily releasable pool of vesicles at hippocampal synapses. Neuron. 16:1197–1207.

Rudolph, S., M.C. Tsai, H. von Gersdorff, and J.I. Wadiche. 2015. The ubiquitous nature of multivesicular release. Trends Neurosci. 38:428–438.

Saitoe, M., T.L. Schwarz, J.A. Umbach, C.B. Gundersen, and Y. Kidokoro. 2001. Absence of junctional glutamate receptor clusters in Drosophila mutants lacking spontaneous transmitter release. Science. 293:514–517.

Salpeter, M.M., and R. Harris. 1983. Distribution and turnover rate of acetylcholine receptors throughout the junction folds at a vertebrate neuromuscular junction. J Cell Biol. 96:1781–1785.

Schotzinger, R.J., and S.C. Landis. 1990. Acquisition of cholinergic and peptidergic properties by sympathetic innervation of rat sweat glands requires interaction with normal target. Neuron. 5:91–100.

Stewart, B.A., H.L. Atwood, J.J. Renmger, J. Wang, and C.F. Wu. 1994. Improved stability of Drosophila larval neuromuscular prepa-rations in haemolymph-like physiological solutions. J Comp Physiol. 175:179–191.

Stuart, G.J., and N. Spruston. 2015. Dendritic integration:60 years of progress. Nat Neurosci. 18:1713–1721.

Sutton, M.A., H.T. Ito, P. Cressy, C. Kempf, J.C. Woo, and E.M. Schuman. 2006. Miniature neurotransmission stabilizes synaptic function via tonic suppression of local dendritic protein synthesis. Cell. 125:785–799.

Tripodi, M., J.F. Evers, A. Mauss, M. Bate, and M. Landgraf. 2008. Structural homeostasis: compensatory adjustments of dendritic arbor geometry in response to variations of synaptic input. PLoS Biol. 6:e260.

Turrigiano, G. 2012. Homeostatic Synaptic Plasticity: Local and Global Mechanisms for Stabilizing Neuronal Function. Cold Spring Harb. Perspect. Biol. 4:a005736.

Turrigiano, G.G. 2008. The self-tuning neuron: synaptic scaling of excitatory synapses. Cell. 135:422–435.

Turrigiano, G.G., and S.B. Nelson. 2004. Homeostatic plasticity in the developing nervous system. Nat Rev Neurosci. 5:97–107.

Vitureira, N., and Y. Goda. 2013. The interplay between Hebbian and homeostatic synaptic plasticity. J Cell Biol. 203:175.

Vitureira, N., M. Letellier, I.J. White, and Y. Goda. 2011. Differential control of presynaptic efficacy by postsynaptic N-cadherin and β-catenin. Nat Neurosci. 15:81–89.

Vukoja, A., U. Rey, A.G. Petzoldt, C. Ott, D. Vollweiter, C. Quentin, D. Puchkov, E. Reynolds, M. Lehmann, S. Hohensee, S. Rosa, R. Lipowsky, S.J. Sigrist, and V. Haucke. 2018. Presynaptic Biogenesis Requires Axonal Transport of Lysosome-Related Vesicles. Neuron. 99:1216–1232 e1217.

Weyhersmuller, A., S. Hallermann, N. Wagner, and J. Eilers. 2011. Rapid active zone remodeling during synaptic plasticity. J Neurosci. 31:6041–6052.

Yuan, Q., Y. Xiang, Z. Yan, C. Han, L.Y. Jan, and Y.N. Jan. 2011. Light-Induced Structural and Functional Plasticity in Drosophila Larval Visual System. Science. 333:1458–1462.

