## Supplementary material for "Distinct target-specific mechanisms homeostatically stabilize transmission at pre-and post-synaptic compartments": Table S1

### Table S1. REAGENTS AND RESOURCES TABLE

| Antibodies | | | |
|---|---|---|---|
| **REAGENT/RESOURCE** | **SOURCE** | **IDENTIFIER** | **DILUTION** |
| Mouse anti-Synapsin (3C11) | Developmental Studies Hybridoma Bank (DSHB) | AB_2313867 | 1:10 |
| Tetramethylrhodamine (TRITC)-conjugated phalloidin (R415) | Thermo Fisher Scientific | 41-6559-05 | 1:1000 |
| Mouse anti-Bruchpilot (nc82) | DSHB | AB_2314866 | 1:100 |
| Guinea pig anti-vGlut | (Goel and Dickman, 2018) | N/A | 1:2000 |
| Mouse anti-GluRIIA (8B4D2) | DSHB | AB_528269 | 1:50 |
| Affinity-Purified Rabbit anti-GluRIIB | (Perry et al., 2017) | N/A | 1:2000 |
| Guinea pig anti-GluRIID | (Kikuma et al., 2017) | N/A | 1:1000 |
| DyLight 405-conjucated secondary antibodies | Jackson ImmunoResearch Laboratories, Inc | 706-475-148 | 1:400 |
| Alexa Fluor 488-conjugated secondary antibodies | Jackson ImmunoResearch Laboratories, Inc | 706-545-148, 715-545-150, 711-545-152 | 1:400 |
| Cy3-conjugated secondary antibodies | Jackson ImmunoResearch Laboratories, Inc | 706-165-148, 715-165-150, 711-165-152 | 1:400 |
| Alexa Fluor 647 conjugated Goat anti-Horseradish Peroxidase | Jackson ImmunoResearch Laboratories, Inc | 123-605-021 | 1:200 |

| Experimental Models: Fly Lines | | |
|---|---|---|
| **REAGENT/RESOURCE** | **REFERENCE** | **SOURCE** |
| *H94-Gal4* | (Davis et al., 1997) | Brain McCabe<br>Ecole polytechnique federale de Lausanne |
| *UAS-FasII-PEST+* | (Davis et al., 1997) | C. Andrew Frank<br>University of Iowa |
| *GluRIIA$^{sp16}$* | (Petersen et al., 1997) | Bloomington Drosophila Stock Center (BDSC #64202) |
| *Cac$^{sfGFP-N}$* | (Gratz et al., 2019) | Kate O'Connor Giles<br>Brown University |
| *UAS-GluRIIA$^{RNAi}$* (p{TRiP.JF02647}attP2) | (Li et al., 2018) | Bloomington Drosophila Stock Center (BDSC #27497) |
| *Tub-FRT-STOP-FRT-Gal4,UAS-FLP,UAS-CD8-GFP* | (Roy et al., 2007) | Brain McCabe<br>Ecole polytechnique federale de Lausanne |

### Supplemental References:


Davis, G.W., C. Schuster, and C.S. Goodman. 1997. Genetic analysis of the molecular mechanisms controlling target selection: target derived Fasciclin II regulates the pattern of synapse formation. *Neuron*. 19:561-573.

Goel, P., and D. Dickman. 2018. Distinct homeostatic modulations stabilize reduced postsynaptic receptivity in response to presynaptic DLK signaling. *Nat Commun*. 9:1856.

Gratz, S.J., P. Goel, J.J. Bruckner, R.X. Hernandez, K. Khateeb, G. Macleod, D. Dickman, and K.M. O'Connor-Giles. 2019. Endogenous tagging reveals differential regulation of Ca2+ channels at single AZs during presynaptic homeostatic potentiation and depression. *J Neurosci*. 39:2416-2429.

Kikuma, K., X. Li, D. Kim, D. Sutter, and D.K. Dickman. 2017. Extended Synaptotagmin Localizes to Presynaptic ER and Promotes Neurotransmission and Synaptic Growth in Drosophila. *Genetics*. 207:993-1007.

Li, X., P. Goel, C. Chen, V. Angajala, X. Chen, and D. Dickman. 2018. Synapse-specific and compartmentalized expression of presynaptic homeostatic potentiation. *Elife*. 7:e34338.

Perry, S., Y. Han, A. Das, and D.K. Dickman. 2017. Homeostatic plasticity can be induced and expressed to restore synaptic strength at neuromuscular junctions undergoing ALS-related degeneration. *Human Mol Genet.* 26:4153-4167.

Petersen, S.A., R.D. Fetter, J.N. Noordermeer, C.S. Goodman, and A. DiAntonio. 1997. Genetic analysis of glutamate receptors in Drosophila reveals a retrograde signal regulating presynaptic transmitter release. *Neuron*. 19:1237-1248.

Roy, B., A.P. Singh, C. Shetty, V. Chaudhary, A. North, M. Landgraf, K. Vijayraghavan, and V. Rodrigues. 2007. Metamorphosis of an identified serotonergic neuron in the Drosophila olfactory system. *Neural Dev* 2:20.
