## Supplementary material for "Distinct target-specific mechanisms homeostatically stabilize transmission at pre-and post-synaptic compartments": Table S2

**Table S2. Absolute values for normalized data and additional statistics.** The figure and panel, genotype, and conditions used are noted. For electrophysiological recordings, average values for mEPSP, EPSP, quantal content (QC), resting membrane potential, input resistance, number of data samples (n), p values, and significance are shown. For confocal imaging analysis, average values for intensity levels of indicated synaptic markers and other relevant parameters such as numbers and area are shown. Standard error values are noted in parentheses. Rows highlighted in blue are the respective controls or baseline values for the particular experiment being referenced.

| **Figure** | **Genotype** | **mEPSP (mV)** | **EPSP (mV)** | **QC** | **mEPSP (Hz)** | **bouton #/muscle** | **QC/ bouton** | **Resting Potential (mV)** | **Input Resistance (MΩ)** | **n** | **P value (significance) (mEPSP amp, EPSP, QC boutons, QC/bouton, mEPSP freq)** |
|---|---|---|---|---|---|---|---|---|---|---|---|
| 1D,E | *$w^{1118}$* (muscle 7) | 0.965 (0.012) | 35.37 (1.99) | 36.65 (2.047) | 2.545 (0.096) | 39.75 (1.509) | 0.922 (0.071) | -68.7 (1.412) | 14.512 (1.216) | 14 | |
| 1D,E | *w;H94-Gal4;UAS-fasII* (muscle 7) | 1.329 (0.063) | 32.08 (1.016) | 24.061 (1.614) | 1.815 (0.155) | 21.81 (2.201) | 1.038 (0.011) | -67.538 (1.409) | 12.013 (0.819) | 16 | 0.0055 (**), >0.9999 (ns), 0.0076 (**), <0.0001 (****), 0.5021 (ns), 0.0091 (**) |
| 1D,E | *$w^{1118}$* (muscle 6) | 1.018 (0.071) | 34.29 (0.815) | 33.683 (1.97) | 2.977 (0.091) | 58.88 (2.856) | 0.572 (0.043) | -64.491 (1.221) | 7.215 (0.237) | 14 | |
| 1D,E | *w;H94-Gal4;UAS-fasII* (muscle 6) | 1.021 (0.015) | 30.84 (0.764) | 30.271 (1.899) | 3.009 (0.148) | 80.14 (1.993) | 0.377 (0.025) | -65.559 (1.720) | 8.790 (0.504) | 16 | >0.9999 (ns), 0.9157 (ns), 0.9667 (ns), 0.0004 (***) 0.0083 (**), 0.9995 (ns) |

| **Figure** | **Genotype** | **Failure rate (0.15 mM $Ca^{2+}$)** | **Paired-pulse facilitation (PPF) rate (0.4 mM $Ca^{2+}$)** | **Paired-pulse depression (PPD) rate (1.5 mM $Ca^{2+}$)** | **n** | **P value (significance) (failure, PPF, PPD)** |
|---|---|---|---|---|---|---|
| 2B,D,F | *$w^{1118}$* (muscle 6) | 57.51 (3.541) | 140.44 (7.128) | 69.55 (4.028) | 9,9,7 | |
| 2B,D,F | *w;H94-Gal4;UAS-fasII* (muscle 6) | 51.02 (5.014) | 167.91 (10.102) | 88.12 (7.662) | 8,9,8 | 0.8813 (ns), 0.0081 (**), 0.0061 (**) |

| **Figure** | **Genotype** | **Estimated mEPSP (mV)** | **Cumulative EPSC (nA)** | **Estimated RRP size** | **n** | **P value (significance) (mEPSC, cum. EPSC, RRP)** |
|---|---|---|---|---|---|---|
| S2C,D | *$w^{1118}$* (muscle 7) | 0.732 (0.027) | 509.88 (40.128) | 695.55 (47.828) | 9 | |
| S2C,D | *w;H94-Gal4;UAS-fasII* (muscle 7) | 0.902 (0.014) | 500.24 (42.102) | 451.29 (40.662) | 8 | 0.0071 (**), 0.8559 (ns), 0.0008 (***) |
| 2G,H | *$w^{1118}$* (muscle 6) | 0.781 (0.026) | 877.97 (56.705) | 1125.55 (105.711) | 9 | |
| 2G,H | *w;H94-Gal4;UAS-fasII* (muscle 6) | 0.827 (0.068) | 1005.21 (63.031) | 1215.67 (93.459) | 8 | 0.6991 (ns), 0.9978 (ns), 0.8761 (ns) |

| **Figure** | **Genotype** | **Functional release site number (N)** | **n** | **P value (significance)** |
|---|---|---|---|---|
| S2G | *$w^{1118}$* (muscle 7) | 468.86 (27.81) | 6 | |
| S2G | *w;H94-Gal4;UAS-fasII* (muscle 7) | 320.3 (11.16) | 6 | 0.0021 (**) |
| 2K | *$w^{1118}$* (muscle 6) | 500.97 (36.22) | 9 | |
| 2K | *w;H94-Gal4;UAS-fasII* (muscle 6) | 410.78 (36.38) | 10 | 0.9711 (ns) |

| Figure | Genotype | BRP puncta #/NMJ | BRP puncta density (#/μm$^2$) | BRP puncta Intensity (% WT) | Total BRP intensity/M7 NMJ (% WT) | n | P value (significance) (BRP puncta #, density, puncta intensity, total intensity) |
|---|---|---|---|---|---|---|---|
| S2B | *w$^{1118}$* (muscle 7) | 141.67 (11.57) | 0.916 (0.014) | 100 (7.526) | 100 (6.664) | 14 | |
| S2B | *w;H94-Gal4;UAS-fasII* (muscle 7) | 99.17 (5.851) | 0.994 (0.111) | 106.821 (8.451) | 78.93 (8.003) | 16 | 0.0049 (**), 0.8976 (ns), 0.6781 (ns), 0.0105 (*) |
| 3D,E,F,G | *w$^{1118}$* (muscle 6) | 261.44 (18.765) | 0.974 (0.087) | 100 (6.167) | 100 (9.778) | 14 | |
| 3D,E,F,G | *w;H94-Gal4;UAS-fasII* (muscle 6) | 309.72 (20.949) | 0.737 (0.062) | 78.76 (7.098) | 94.77 (9.175) | 16 | 0.0205 (*), 0.0051 (**), 0.0084 (**), 0.8846 (ns) |

| Figure | Genotype | Cac puncta #/NMJ | Cac puncta density (#/μm$^2$) | Cac puncta intensity (% WT) | Total Cac intensity/M7 NMJ (% WT) | n | P value (significance) (Cac puncta #, density, puncta intensity, total intensity) |
|---|---|---|---|---|---|---|---|
| 3D,E,F,G | *cac$^{sfGFP-N}$*(muscle 6) | 132.44 (10.628) | 0.881 (0.064) | 100 (7.443) | 100 (9.803) | 13 | |
| 3D,E,F,G | *cac$^{sfGFP-N}$; H94-Gal4;UAS-fasII* (muscle 6) | 141.68 (11.917) | 0.629 (0.053) | 81.97 (7.255) | 92.88 (8.765) | 12 | 0.6267 (ns), 0.0053 (**), 0.0075 (**), 0.9121 (ns) |

| Figure | Genotype | GluRIIA puncta #/NMJ | GluRIIB puncta #/NMJ | GluRIID puncta #/NMJ | n | P value (significance) (GluRIIA, GluRIIB, GluRIID) |
|---|---|---|---|---|---|---|
| S1C | *w$^{1118}$* (muscle 7) | 164.63 (9.977) | 155.53 (10.093) | 168.21 (11.288) | 9 | |
| S1C | *w;H94-Gal4;UAS-fasII* (muscle 7) | 92.9 (6.163) | 85.133 (6.292) | 72.66 (9.035) | 10 | 0.0007 (***), 0.0005 (***), 0.0003 (***) |
| S1G | *w$^{1118}$* (muscle 6) | 208.125 (14.109) | 218.33 (13.751) | 197.75 (13.607) | 9 | |
| S1G | *w;H94-Gal4;UAS-fasII* (muscle 6) | 278.11 (20.109) | 289.71 (22.892) | 267.516 (19.955) | 10 | 0.0049 (**), 0.0068 (**), 0.0071 (**) |

| Figure | Genotype | GluRIIA puncta intensity (%WT) | GluRIIB puncta intensity (%WT) | GluRIID puncta intensity (%WT) | n | P value (significance) (GluRIIA, GluRIIB, GluRIID) |
|---|---|---|---|---|---|---|
| S1D | *w$^{1118}$* (muscle 7) | 100 (6.271) | 100 (5.828) | 100 (7.188) | 9 | |
| S1D | *w;H94-Gal4;UAS-fasII* (muscle 7) | 162.54 (11.11) | 155.21 (12.292) | 151.09 (14.65) | 10 | 0.0068 (**), 0.0059 (**), 0.0079 (**) |
| S1H | *w$^{1118}$* (muscle 6) | 100 (4.769) | 100 (6.715) | 100 (5.407) | 9 | |
| S1H | *w;H94-Gal4;UAS-fasII* (muscle 6) | 102.54 (10.98) | 95.21 (13.03) | 91.12 (13.955) | 10 | 0.6991 (ns), 0.9978 (ns), 0.8761 (ns) |

| Figure | Genotype | mEPSP (mV) | EPSP (mV) | QC | GluRIIA intensity (%WT) | bouton #/muscle | Resting Potential (mV) | Input Resistance (MΩ) | n | P value (significance) (mEPSP, EPSP, QC, GluRIIA, bouton #) |
|---|---|---|---|---|---|---|---|---|---|---|
| 4E | $w^{1118}$ (muscle 7) | 0.96 (0.012) | 35.37 (1.99) | 36.65 (2.047) | 100 (4.961) | 39.75 (1.509) | -68.7 (1.412) | 14.512 (1.216) | 14 | |
| 4E | *w;Tub-FRT-STOP-FRT-Gal4,UAS-FLP,UAS-CD8-GFP;H94-Gal4,nSyb-Gal80/UAS-FasII* (muscle 7) | 1.369 (0.041) | 33.19 (1.536) | 24.24 (1.614) | 168.15 (16.053) | 18.75 (2.201) | -63.713 (2.009) | 12.013 (0.819) | 12 | 0.0002 (***), 0.9885 (ns), 0.0045 (**), 0.0041 (**) <0.0.0001 (****) [Compared to WT-muscle7] |
| 4E | *w;Tub-FRT-STOP-FRT-Gal4,UAS-FLP,UAS-CD8-GFP;H94-Gal4,nSyb-Gal80/UAS-FasII;UAS-GluRIIA*$^{RNAi}$ (muscle 7) | 1.35 (0.039) | 31.71 (1.991) | 22.37 (1.211) | 156.74 (21.191) | 19.21 (0.945) | -65.51 (1.875) | 15.51 (0.98) | 11 | >0.9999 (ns), >0.9999 (ns), 0.9976(ns), >0.9999 (ns), >0.9999 (ns) |
| 4F | $w^{1118}$ (muscle 6) | 1.018 (0.071) | 34.29 (0.815) | 33.683 (1.97) | 100 (5.543) | 58.88 (2.856) | -64.491 (1.221) | 7.215 (0.237) | 14 | |
| 4F | *w;Tub-FRT-STOP-FRT-Gal4,UAS-FLP,UAS-CD8-GFP;H94-Gal4,nSyb-Gal80/UAS-FasII* (muscle 6) | 0.97 (0.022) | 32.94 (0.894) | 33.95 (1.659) | 100 (6.78) | 72.44 (5.551) | -65.559 (1.720) | 9.120 (0.414) | 12 | 0.9786 (ns), >0.9999 (ns), >0.9999 (ns), 0.9997 (ns), 0.0045 (**) [Compared to WT-muscle6] |
| 4F | *w;Tub-FRT-STOP-FRT-Gal4,UAS-FLP,UAS-CD8-GFP;H94-Gal4,nSyb-Gal80/UAS-FasII;UAS-GluRIIA*$^{RNAi}$ (muscle 6) | 0.46 (0.027) | 31.94 (1.708) | 66.54 (3.732) | 10.79 (2.231) | 70.99 (5.96) | -62.87 (1.591) | 11.69 (1.065) | 11 | <0.0001 (****), 0.9996 (ns), <0.0001 (****), <0.0001 (****),>0.9999 (ns) |

| Figure | Genotype | PhTx | mEPSP (mV) | EPSP (mV) | QC | Resting Potential (mV) | Input Resistance (MΩ) | n | P value (significance) (mEPSP, EPSP, QC) |
|---|---|---|---|---|---|---|---|---|---|
| S3B | $w^{1118}$ (muscle 7) | - | 0.954 (0.028) | 33.1 (1.547) | 34.59 (2.292) | -64.31 (2.015) | 12.35 (0.993) | 9 | |
| S3B | $w^{1118}$ (muscle 7) | + | 0.5 (0.026) | 31.11 (2.15) | 62.22 (4.193) | -66.55 (3.036) | 10.95 (0.768) | 9 | <0.0001 (****), 0.9995 (ns), <0.0001 (****) |
| S3C | *w;Tub-FRT-STOP-FRT-Gal4,UAS-FLP,UAS-CD8-GFP;H94-Gal4,nSyb-Gal80/UAS-FasII;UAS-GluRIIA*$^{RNAi}$ (muscle 7) | - | 1.35 (0.039) | 31.71 (1.991) | 22.37 (1.211) | -65.51 (1.875) | 15.51 (0.98) | 11 | |
| S3C | *w;Tub-FRT-STOP-FRT-Gal4,UAS-FLP,UAS-CD8-GFP;H94-Gal4,nSyb-Gal80/UAS-FasII;UAS-GluRIIA*$^{RNAi}$ (muscle 7) | + | 0.51 (0.026) | 28.02 (1.424) | 54.94 (3.767) | -67.79 (2.891) | 13.39 (1.554) | 10 | <0.0001 (****), 0.9982 (ns), <0.0001 (****) |
| S3D | $w^{1118}$ (muscle 6) | - | 0.94 (0.022) | 34.98 (1.076) | 37.21 (1.981) | -67.09 (2.296) | 6.95 (0.316) | 9 | |
| S3D | $w^{1118}$ (muscle 6) | + | 0.45 (0.019) | 31.83 (0.873) | 70.33 (4.137) | -63.94 (2.095) | 7.21 (0.591) | 9 | <0.0001 (****), 0.9957 (ns), <0.0001 (****) |
| S3E | *w;Tub-FRT-STOP-FRT-Gal4,UAS-FLP,UAS-CD8-GFP;H94-Gal4,nSyb-Gal80/UAS-FasII;UAS-GluRIIA*$^{RNAi}$ (muscle 6) | - | 0.46 (0.027) | 31.94 (1.708) | 66.54 (3.732) | -62.87 (1.591) | 11.69 (1.065) | 11 | |
| S3E | *w;Tub-FRT-STOP-FRT-Gal4,UAS-FLP,UAS-CD8-GFP;H94-Gal4,nSyb-Gal80/UAS-FasII;UAS-GluRIIA*$^{RNAi}$ (muscle 6) | + | 0.37 (0.012) | 29.38 (0.989) | 79.5 (4.575) | -64.48 (1.592) | 9.09 (0.855) | 10 | 0.2349 (ns), >0.9999 (ns), 0.3159 (ns) |
