## Supplementary figures and images for "Distinct target-specific mechanisms homeostatically stabilize transmission at pre-and post-synaptic compartments"

### Supplemental Figures 1-3

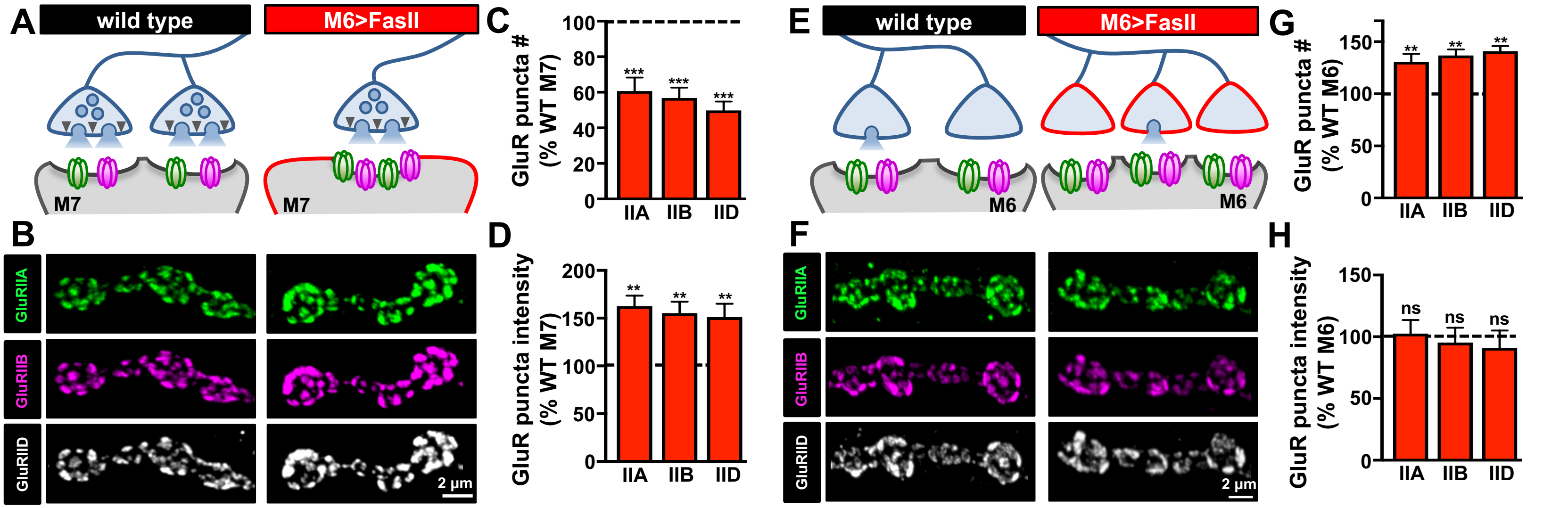

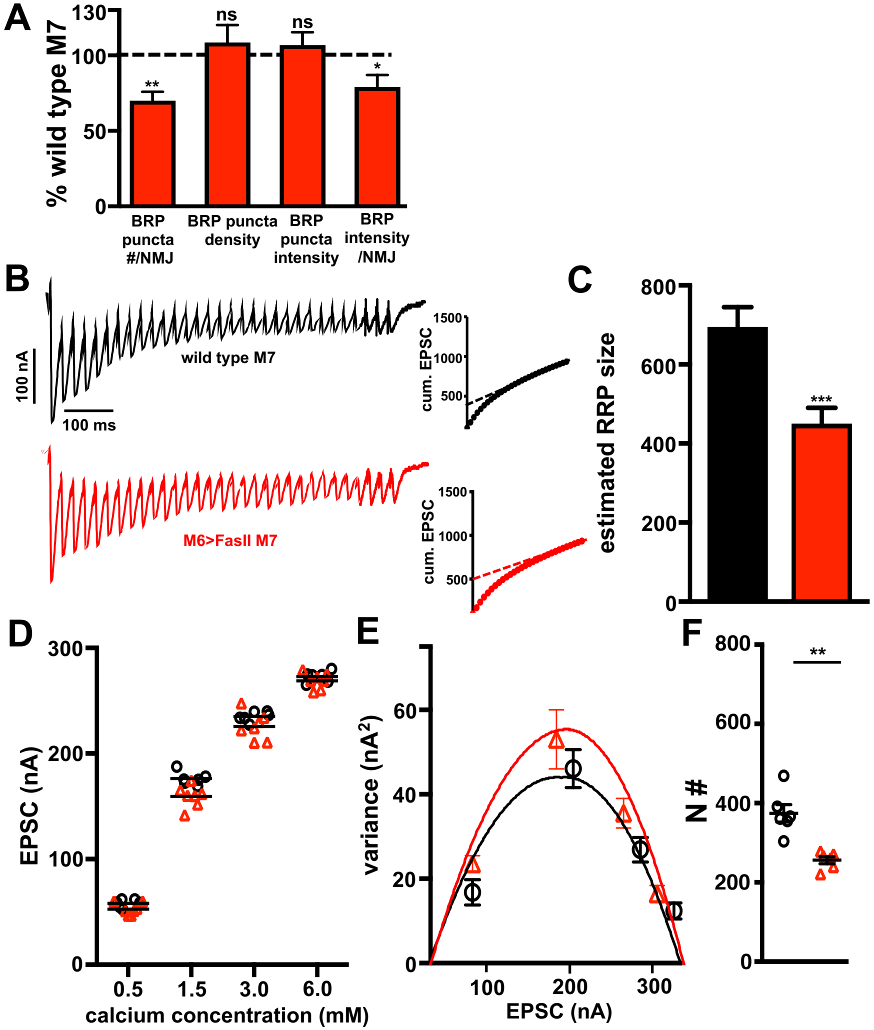

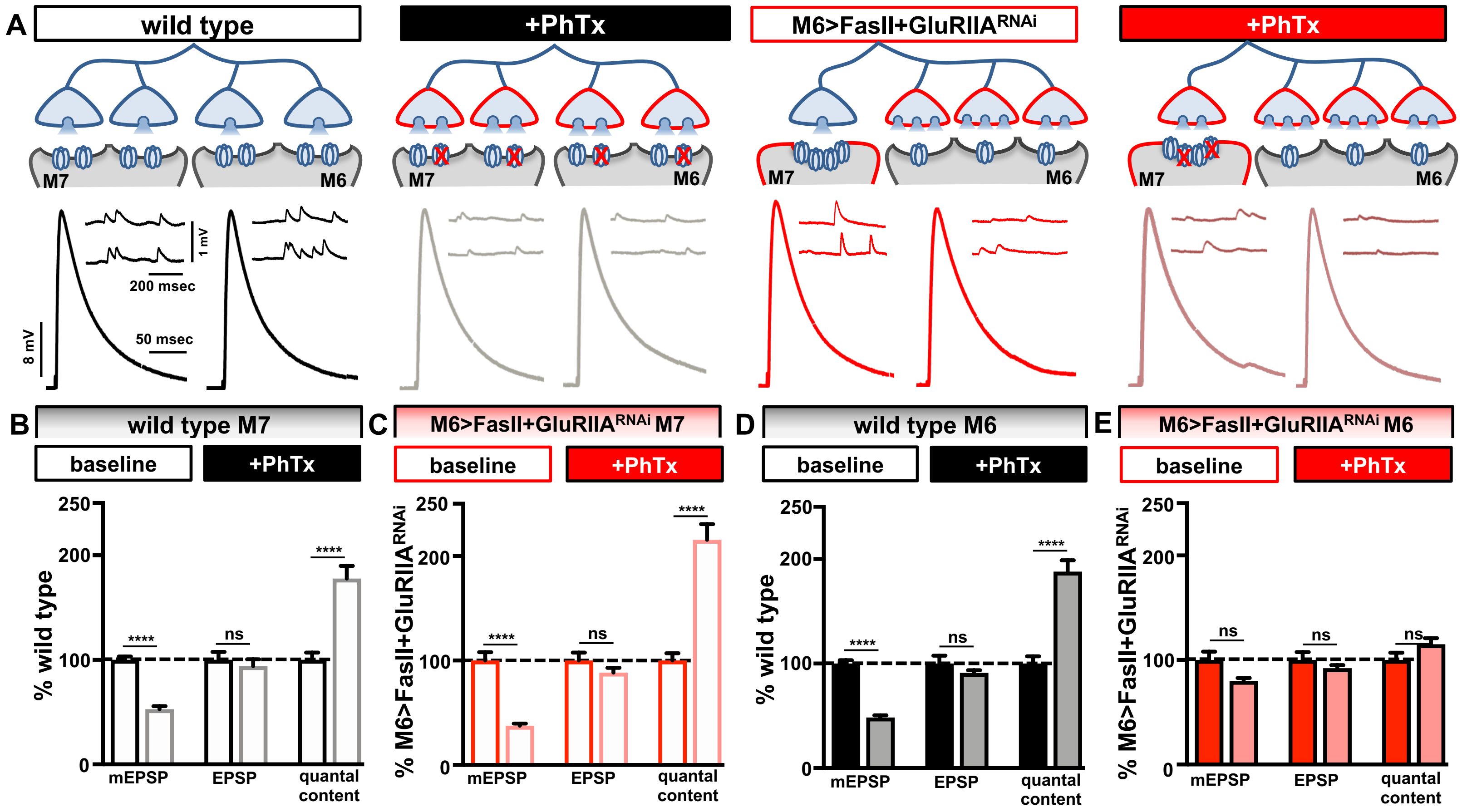
